# ggalign: Bridging the Grammar of Graphics and Biological Multilayered Complexity

**DOI:** 10.1101/2025.02.06.636847

**Authors:** Yun Peng, Yuxuan Song, Peng Luo, Jianfeng Li, Jian-Guo Zhou, Guangchuang Yu, Tao Xu, Shixiang Wang

## Abstract

Composable visualization is essential in biomedical research, particularly when working with complex, multi-dimensional biological data. However, many existing tools focus on specific fields and lack the comprehensive features needed to capture the intricacies and interrelations of datasets. To address this, we introduce ggalign, an R package built on the grammar of graphics framework, providing an easy-to-use platform for creating composable visualizations of interconnected data. It offers a variety of layout options, including *StackLayout, QuadLayout*, and *CircleLayout*, tailored to different data types, and supports flexible multi-dimensional relationships, such as one-to-many, many-to-one, many-to-many, and crosswise connections. Additionally, ggalign enhances the clarity of visualizations through its annotation features. Overall, ggalign offers a powerful and flexible solution for visualizing complex data in biomedical research.

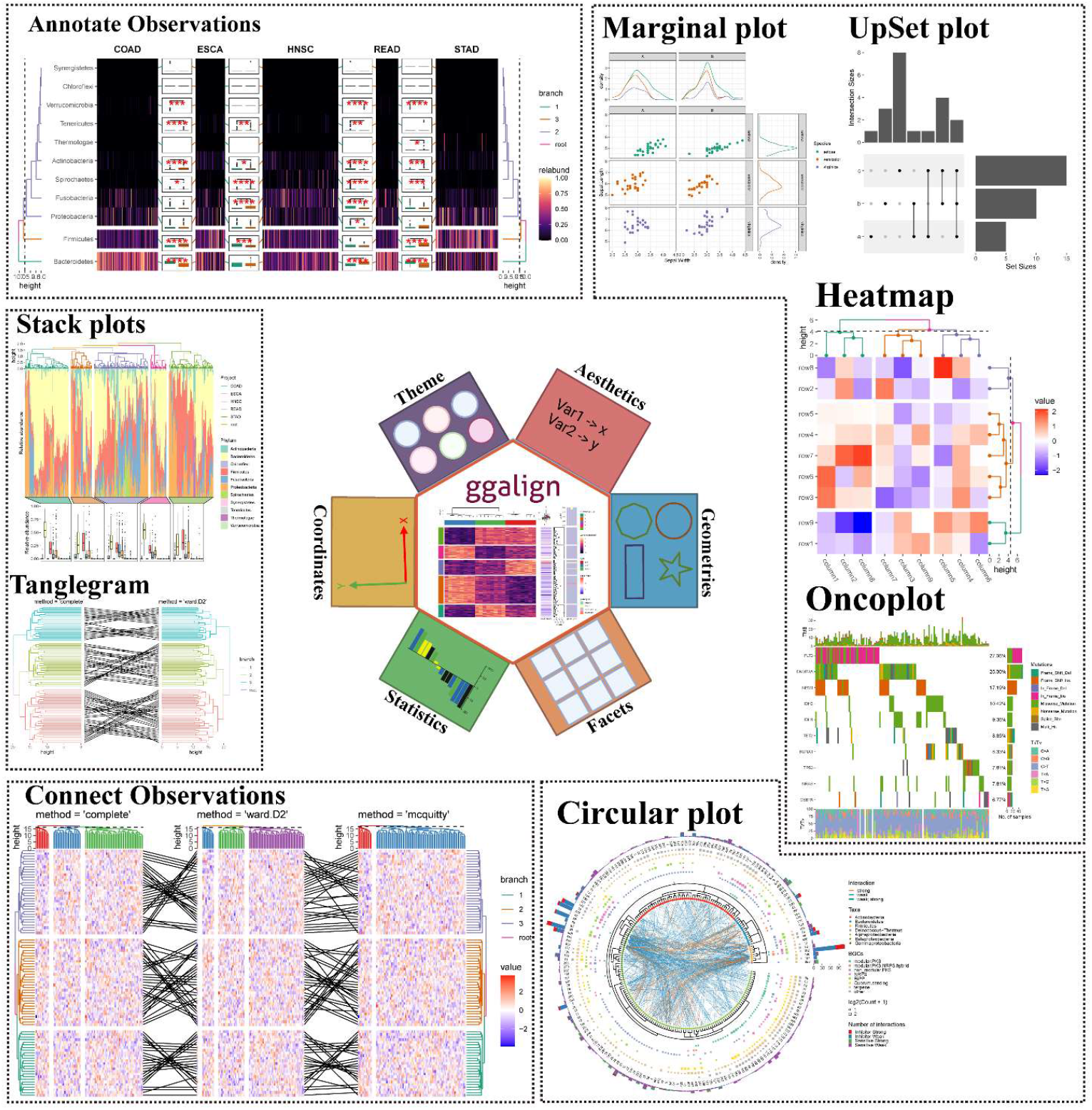

## Introduction

Data visualization is a cornerstone of modern data analysis, enabling the effective exploration, interpretation, and communication of complex datasets. The integration and visualization of multi-dimensional biological data has become a critical challenge in modern biomedical research. With the advent of high-throughput technologies, datasets now span multiple layers, such as genomic, transcriptomic, epigenomic, and microbiome data, requiring tools that can effectively organize and present this complexity, and composable visualization has been extensively used. In this context, composable visualization and becomes increasingly important, as it enables flexible and dynamic representation of data across various dimensions, allowing researchers to uncover intricate relationships and patterns.

While several specialized packages exist to support the creation of specific visualizations, such as ComplexHeatmap^1^ for heatmaps and ggtree^2^ for phylogenetic trees, these tools are often difficult to adapt for use with other plot types. Tools like aplot^3^ and marsilea^4^ provide excellent composable visualizations, but are limited to one-to-one relationship alignment. This restriction makes them less suitable for handling multi-layered, interconnected omics datasets, which frequently involve one-to-many or many-to-one mappings, such as between different taxonomic levels or between pathways and genes.

The ggalign package bridges this gap by providing a unified framework for multi-plot visualization based on the grammar of graphics. By integrating the powerful visualization capabilities of ggplot2 with well-designed advanced linking features, ggalign make it possible to produce publication-ready visualizations that are both aesthetically pleasing and scientifically accurate. This article presents an overview of the ggalign package, exploring its design, core functionalities, and the specific linking features it provides. By offering a robust solution to the challenges of multi-plot layout, ggalign enhances the ability to generate high-quality, composable visualizations that improve the interpretability and communication of complex data insights.

## Results

### Design philosophy

The ggalign helps unlock insights from data by making it easier to structure, align, and analyze various datasets. It is built on the foundational principles of the grammar of graphics, providing an advanced, object-oriented approach to multi-plot visualization. At the core of its design is the layout system that facilitates the intuitive arrangement of plots. This system offers three primary *Layout* classes, each tailored to different visualization needs, enabling the alignment of both discrete and continuous variables **(Figure S1 and Figure 1)**: *StackLayout*: This class arranges plots in a stacked format, either horizontally or vertically **(Figure S1A)** (Figure 1). It is ideal for comparing datasets or dimensions within a single dataset that share similar axes or data types. A variation of this layout, *StackCross*, allows for linking two plots with different numbers of observations, orderings, or groups. This makes it suitable for visualizing datasets with differing structures while maintaining clarity and consistency. *QuadLayout*: This class arranges plots in four quadrants around a central plot, with the central plot acting as a reference point **(Figure S1B)**. This layout offers additional context by placing surrounding plots in a way that complements or enriches the main plot’s information. A specific variant, *HeatmapLayout*, simplifies the creation of heatmap plots by automatically applying the necessary data mappings and visualization layers. *CircleLayout*: This class arranges plots in a circular pattern, where each plot forms a circular track **(Figure S1C)**. This layout offers a dynamic and visually engaging alternative to traditional grid-based layouts, making it especially suitable for representing cyclical or interconnected relationships within data. By using ggalign, researchers can unlock new insights from their data by organizing it in ways that enhance pattern recognition, simplify complex relationships, and reveal hidden trends, all of which are critical for advancing research and data-driven discoveries.

**Figure 1:**
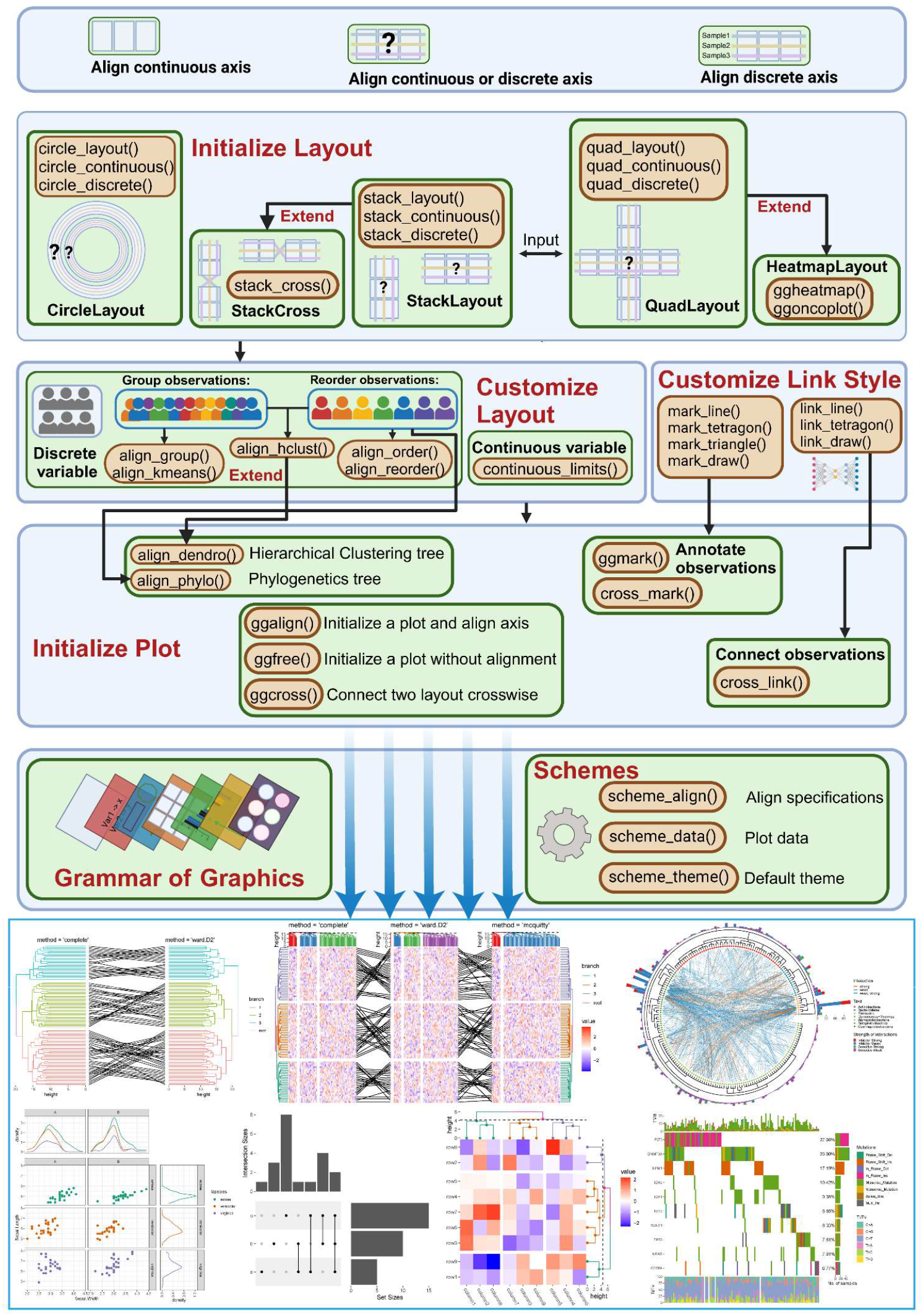
ggalign design and workflow. The ggalign process begins by initializing the layout and choosing the appropriate layout based on the nature of your data and the comparisons you wish to make. Next, for continuous axes, you can use *continuous_limits()* to set the axis ranges. For discrete axes, you can reorder the data points using *align_order()* or *align_reorder()*, or split them into different groups with *align_group()* or *align_kmeans()*. Additionally, you can reorder or group data simultaneously based on hierarchical clustering with *align_hclust()*. After customizing the layout, users can add plots with *ggalign(), ggfree()*, or *ggcross()*, depending on whether they want to align the axes or connect two plots crosswise. The functions *align_dendrogram()* and *align_phylo()* allow users to integrate hierarchical or phylogenetic tree-like structures into the layout, both function will also customize the layout based on the tree structure. Users can also use *ggmark()* or *cross_mark()* to add plots for annotating selected observations, or use *cross_link()* to connect observations between two different plots. Additional ggplot2 elements such as geoms, stats, or scales can be layered to refine the visualization. Simultaneously, *scheme_align()* can be used to control the final assembly specifications, such as how to collect guide legends, *scheme_data()* to transform the plot data, and *scheme_theme()* to set the default theme.

**Figure S1:**
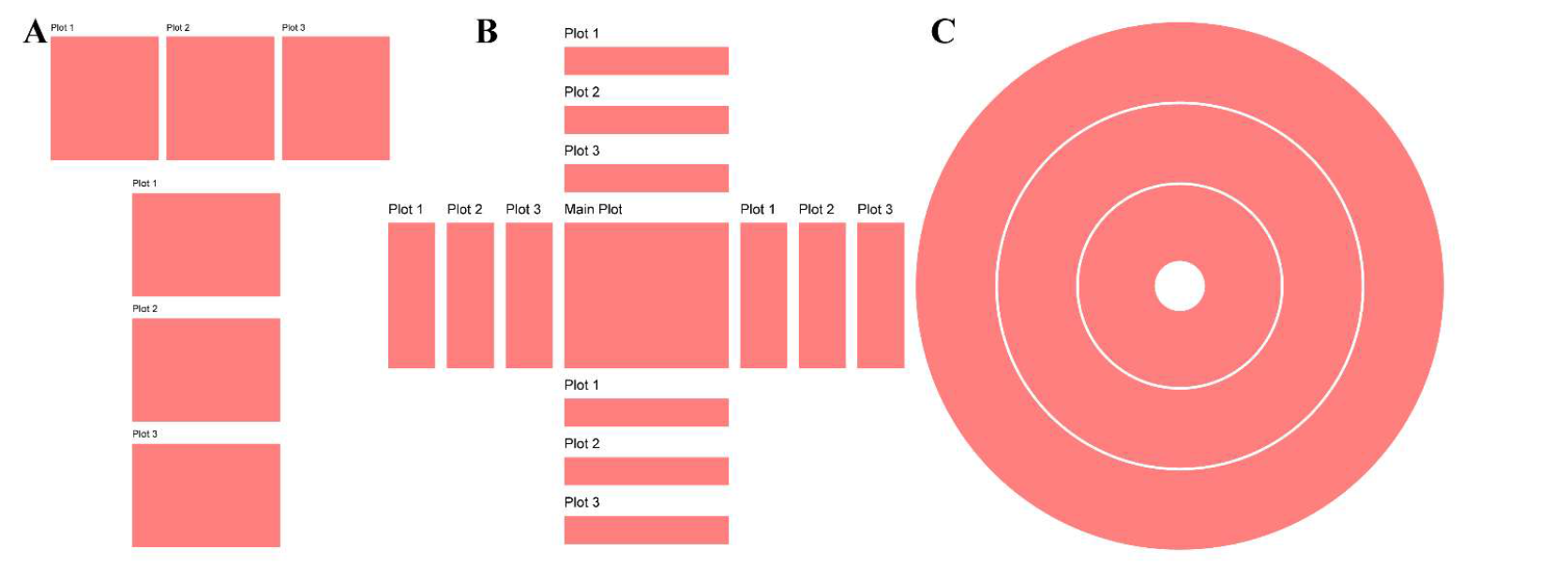
Layout Classes. **(A)** StackLayout for both horizontal and vertical **(B)** QuadLayout **(C)** CircleLayout

Beyond these layouts, ggalign introduces advanced linking features for handling discrete axes, enabling the connection of plots in one-to-many, many-to-one, many-to-many, and crosswise relationships (**Figure 2A-D**). This flexibility enhances the ability to compare and align plots with varying structures. The linking feature also allows for the annotation of specific subsets of observations, thereby improving both the clarity and interactivity of the visualizations **(Figure 3)**.

**Figure 2:**
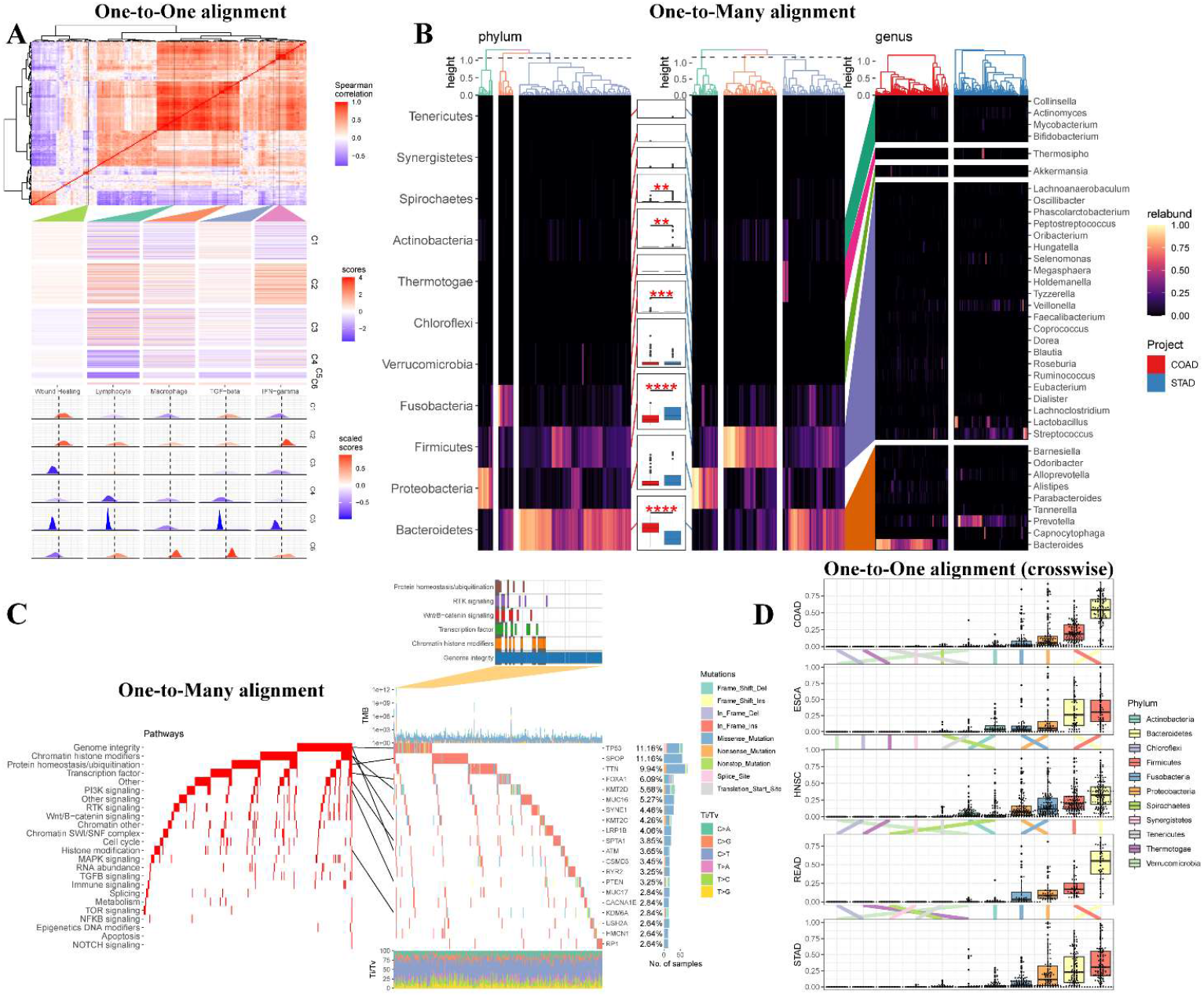
Visualizing complex data relationships with ggalign. **(A)** Expression signature modules and identification of immune subtypes. Five representative gene expression signatures (linked with colored polygon) were used to cluster TGCA tumor samples (rows), resulting in 6 immune subtypes C1–C6. Then the distributions of signature scores within the 6 subtypes (rows) were depicted, with dashed line indicating the median. **(B)** Differential phyla and corresponding differential genera between COAD and STAD were linked with colored polygon. Label “*” means P < 0.05, label “**” means P <0.01, label “***” means P < 0.001, and label “****” means P < 0.0001. **(C)** Oncoprints of both pathways and gene mutations were depicted, with the lines connecting specific genetic alterations. The most significantly involved pathways in TP53-mutated patients were annotated. **(D)** The transition of phylum ordering based on the mean relative abundance across the 6 tumor types was shown with colored lines.

**Figure 3:**
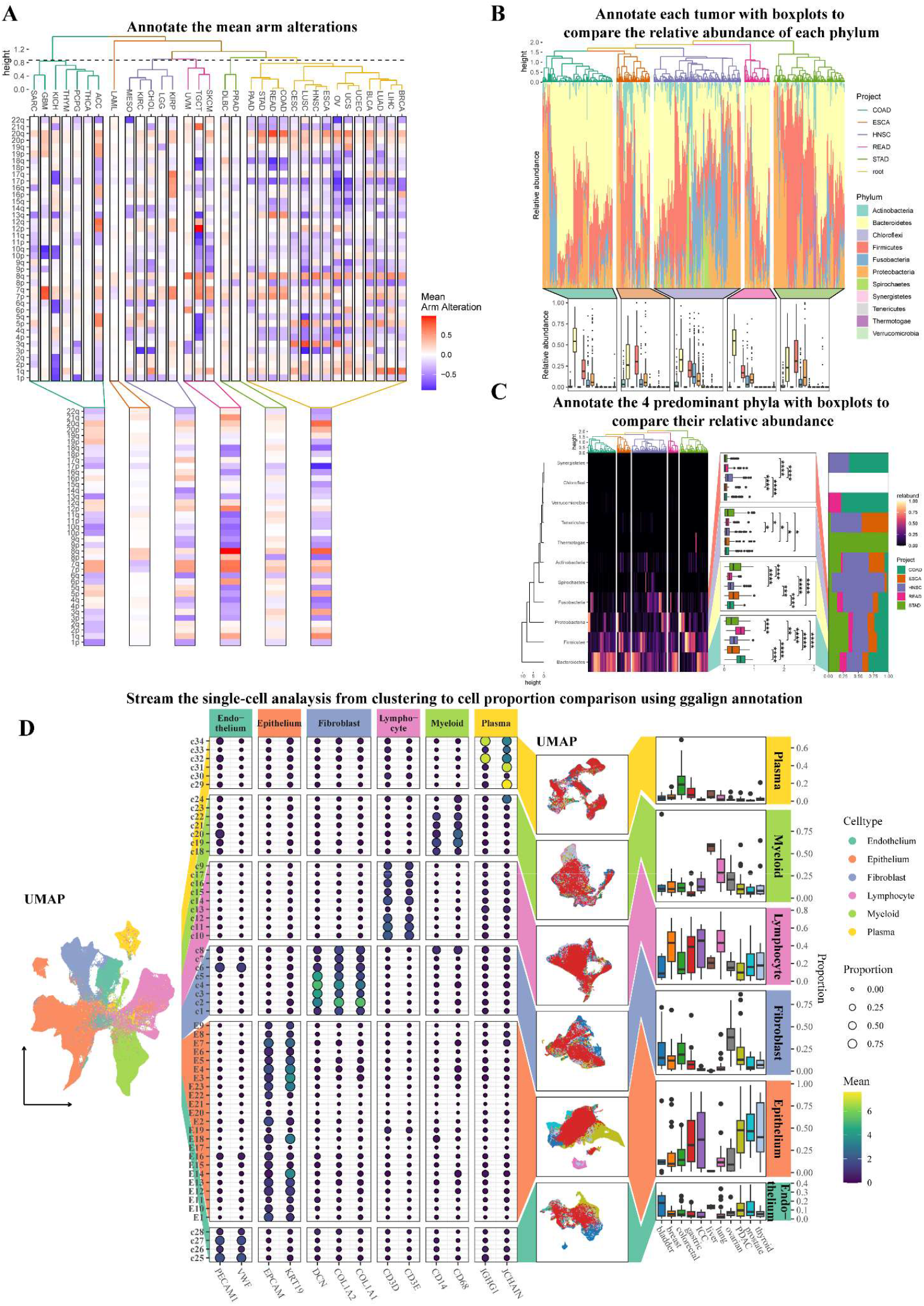
Comprehensive annotation of data patterns with ggalign. **(A)** The arm-level alterations were depicted and hierarchical clustering using the ward.D2 agglomeration method grouped the tumor types into 6 distinct clusters, with the mean arm alterations annotated. **(B)** The relative abundance of the intratumor microbiome at the phylum level across 5 tumor types from TCGA was depicted. A boxplot was used to compare the relative abundance across different phyla. **(C)** The relative abundance of the intratumor microbiome at the phylum level across 5 tumor types from TCGA was depicted. The top 4 phyla (“Bacteroidetes”, “Firmicutes”, “Proteobacteria”, and “Fusobacteria”) were selected, and a boxplot was used to compare their abundance across the different tumor types. Label “*” means P < 0.05, label “**” means P <0.01, label “***” means P < 0.001, and label “****” means P < 0.0001. **(D)** From left to right. The first plot depicted the UMAP plot of all cells. Canonical cell markers were then used to annotate the clusters, identifying 6 distinct cell types. Each cell type was visualized using a separate UMAP plot to compare its distribution across tumor types. Boxplots were used to assess the differences in cell proportions across these tumor types.

While other composable visualization tools, such as aplot^3^ and marsilea^4^, primarily support one-to-one relationship and offer limited layout options **(Table 1)**, ggalign provides a unified and versatile framework for organizing and visualizing complex data, offering extensive customization options. In addition to the commonly used *StackLayout* and *QuadLayout*, ggalign also introduces *CircleLayout*, further enhancing its ability to structure and align data for comprehensive visual exploration **(Figure S1C and Figure S2)**. Additionally, ggalign enhances the exploration and mining of data by allowing users to seamlessly integrate geometric layers from the extensive ggplot2 ecosystem. Specialized tools like ComplexHeatmap are optimized for specific visualization types, such as heatmaps, but are often constrained to particular use cases. In contrast, ggalign supports a broad range of visualization types, offering a more generalized and adaptable solution suitable for diverse research contexts **(Table 1)**. A detailed comparison of other specifications with these tools can be found in **Table S1**. Its flexibility makes it especially valuable in fields like omics research, where the organization and visualization of complex, associated data are crucial for comprehensive analysis and discovery.

**Table 1:** Comparison with other similar tools.

|  |  |  |  |  |  |
| --- | --- | --- | --- | --- | --- |
|  |  | ggalign | marsilea | aplot | ComplexHeatmap |
| Language |  | R | Python | R | R |
| User Interface |  | Declarative | Declarative | Declarative+Functional | Functional |
| Plot System |  | ggplot2<br>(Advanced plot system built on grid system) | Matplotlib | ggplot2 (Advanced plot system built on grid system) | grid |
| Focus |  | Composable Visualization | Composable Visualization | Composable Visualization | Heatmap |
| StackLayout |  | √ | √ | √ | √ |
| QuadLayout |  | √ | √ | √ | Heatmap only<br>(discrete variables) |
| CircleLayout |  | √ | × | × | × |
| Relationship | One-to-One | √ | √ | √ | √ |
|  | One-to-Many/<br>Many-to-One | √ | × | × | × |
|  | Many-to-Many | √ | × | × | × |
|  | Crosswise | √ | × | × | × |
| Annotate observations |  | √ | × | × | √ |
| Compatible with ggplot2 |  | √ | × | √ | × |

**Figure S2:**
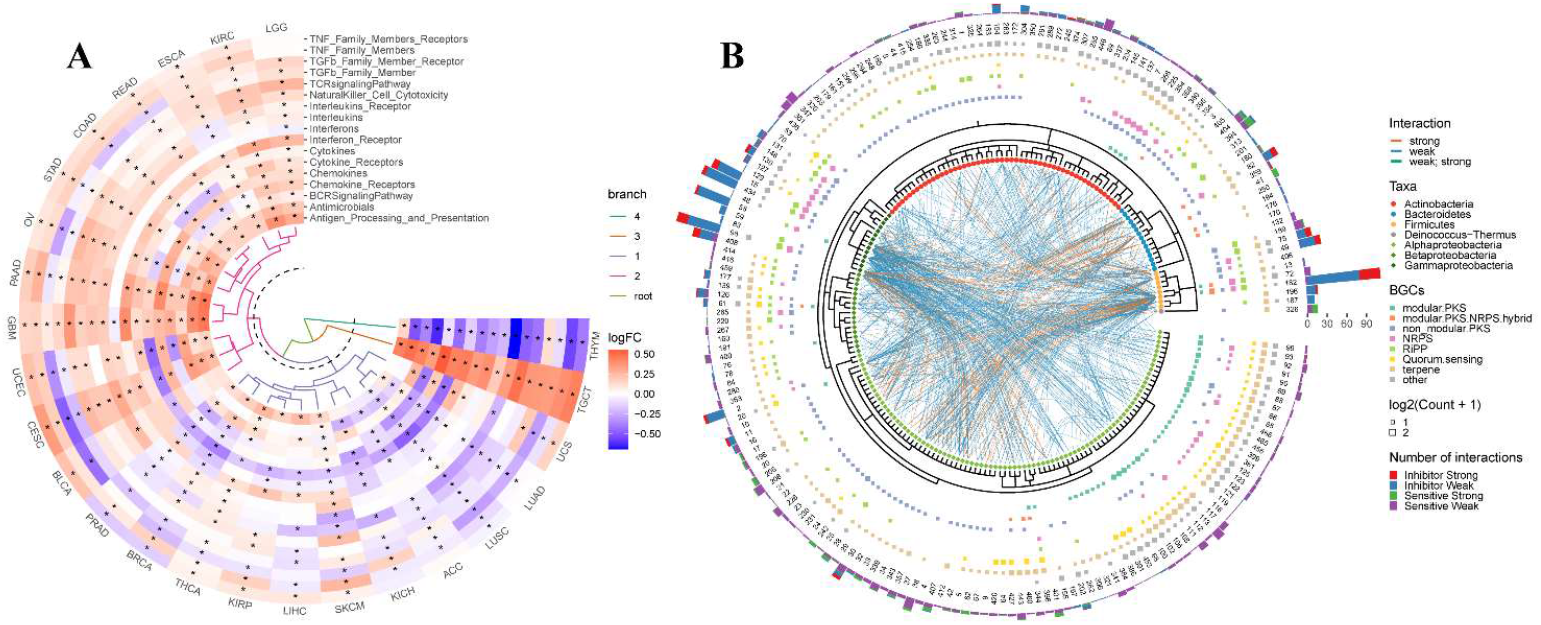
Examples of CircleLayout. (A) Circular Heatmap displaying the comparison of immune-related pathway expression levels between tumor and normal samples across 25 cancer types. Significant dysregulations are marked with an asterisk (“*”). (B) circular phylogenetic tree combined with a chord diagram to represent multidimensional datasets and the inter-relationships between different biological elements.

**Table S1:** Comparison of specifications with other similar tools.

| Specification | ggalign | marsilea | aplot | ComplexHeatmap |
| --- | --- | --- | --- | --- |
| Reorder observations | √ | √ | × | Heatmap Only |
| Group observations into different panels | √ | √ | × | Heatmap Only |
| Clustering algorithm | Kmeans,Hierarchical Clustering and arbitrary algorithm | × | × | Kmeans,Hierarchical Clustering and arbitrary algorithm |
| Legends Creation | Automatic | Manual | Automatic | Limited automatic, requires manual add |
| Legends Position | Anywhere, can be controlled for a single plot | Anywhere | Anywhere | Four sides, can only be placed on one side at a time |
| Dendrogram | Tree from both hclust or ape | hclust only | Tree from both hclust or ape | hclust only |
| Tanglegram | √ | × | × | × |
| 3D Heatmap | √ | × | × | √ |
| Oncoplot | √ | √ | √ | √ |
| UpSet plot | √ | √ | × | √ |

### ggalign workflow

The workflow for using ggalign is intuitive and follows a style similar to ggplot2, utilizing the “+” operator **(Figure 1 and Figure S3)**. The process begins by initializing the layout and selecting the appropriate layout based on the nature of your data and the comparisons you intend to make. After setting up the layout, you can customize it further and initialize the plot. Additional ggplot2 elements, such as geoms, stats, or scales, can be layered to refine the visualization. Simultaneously, schemes can be applied to control the final assembly specifications, manage plot data, and set the default theme. Finally, printing the layout generates the complete visualization.

**Figure S3:**
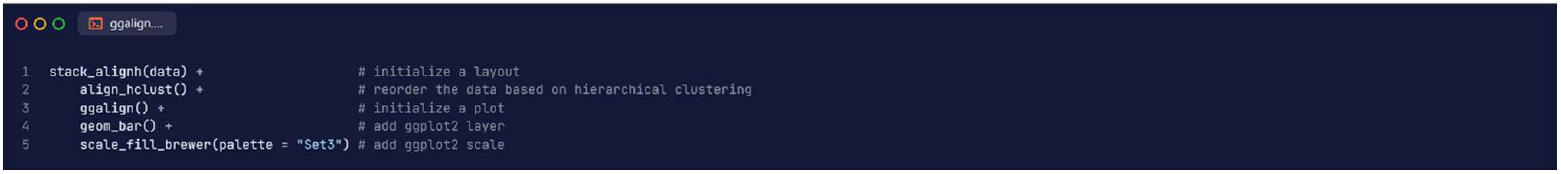
Example code for ggalign workflow.

We showcase ggalign functionalities by applying it to several published datasets, including microbiome research, single-cell research, transcriptomics, and genomics research.

### Link for complex relationship data analysis

The ggalign package offers enhanced one-to-one alignment features, by allowing users to link specific subsets of observations. By focusing on these subsets, users can ensure that the visualized results are both informative and relevant, highlighting key patterns or trends without being overwhelmed by extraneous information. Using ggalign, we mimicked the figure from Thorsson et al.^5^, where 5 representative immune signatures were aligned to cluster TCGA tumor samples, resulting in 6 distinct immune subtypes (C1–C6) **(Figure 2A)**.

Additionally, the ggalign package can handle one-to-many, many-to-one, and even many-to-many relationship data, which are particularly useful for explaining complex data or presenting summarized statistics. These features are especially valuable in omics research, where inconsistent mapping often occurs across different data types, such as pathways to genes, genes to transcripts, genes to methylation probes, or between different taxonomic levels. This capability facilitates clearer comparisons and deeper insights across diverse biological data layers. To demonstrate this feature, we use the ggalign to explore the differential phyla and their corresponding genera between colon adenocarcinoma (COAD) and stomach adenocarcinoma (STAD) **(Figure 2B)**. The ggalign helps to seamlessly link the phyla with their genera (one-to-many relationship), allowing for a clearer visualization of the relationships. Additionally, ggalign is used to generate oncoprints of both pathways and gene mutations in prostate cancer (PRAD), highlighting the interactions between pathways and genes **(Figure 2C)**. The flexibility of ggalign’s linking features ensures that complex relationships in the data are presented in an organized and interpretable manner.

Furthermore, ggalign allows for crosswise alignment, which is especially useful for visualizing transitions and enhancing our ability to observe trends and patterns. For example, we employed crosswise alignment to visualize the transition in the ordering of phyla based on their mean relative abundance across the 5 tumor types **(Figure 2D)**, which revealed that the top 4 phyla were consistent, contributing the majority of the microbiome abundance compared to other phyla.

### Enhanced annotating function for data interpretation

The enhanced annotating function provided by ggalign allows for effective annotation, significantly improving both the clarity and interactivity of the visualizations. By simplifying the process of adding relevant annotations, ggalign ensures that important patterns and insights are clearly highlighted, making complex data easier to interpret and explore. For example, we used ggalign to visualize arm-level alterations and performed hierarchical clustering using the ward.D2 agglomeration method. This method grouped the tumor types into 6 distinct clusters, and ggalign allowed us to annotate the mean arm alterations for each group in a clear and informative manner **(Figure 3A)**. In another example, ggalign was employed to examine the relative abundance of the intratumor microbiome at the phylum level across 5 tumor types, with a boxplot to compare the relative abundance across different phyla within each tumor type **(Figure 3B)**. Similarly, ggalign was used to generate a heatmap displaying the relative abundance of each phylum across the five tumor types, with boxplots to highlight the comparison of 4 predominant phyla (“Bacteroidetes,” “Firmicutes,” “Proteobacteria,” and “Fusobacteria”) across tumor types **(Figure 3C)**. The effective annotating capabilities ensured that complex patterns were presented in an intuitive manner, facilitating more insightful data exploration and analysis. Using ggalign’s annotation features, we can also streamline our analysis process effectively. Take single-cell analysis as an example **(Figure 3D)**: to compare the proportions of different cell types, we first clustered the cells and visualized the results with a Uniform Manifold Approximation and Projection (UMAP)plot. Canonical cell markers were then used to annotate the clusters, allowing us to identify 6 distinct cell types: epithelium (*EPCAM* and *KRT19*), lymphocytes (*CD3D* and *CD3E*), myeloid cells (*CD14* and *CD68*), fibroblasts (*DCN, COL1A2*, and *COL1A1*), endothelium (*PECAM1* and *VWF*), and plasma cells (*IGHG1* and *JCHAIN*). Each cell type was visualized with a separate UMAP plot to compare its distribution across tumor types. Finally, boxplots were used to assess differences in cell proportions across these tumor types. By providing intuitive and informative annotations, ggalign improves the presentation of complex data, and facilitates insightful data exploration, making it a powerful tool for various research applications.

## Discussion

In this study, we introduced ggalign, a versatile and composable visualization tool designed to address the growing complexity of multi-dimensional biological datasets in biomedical research. By leveraging the powerful framework of ggplot2, ggalign provides a unified platform for the creation of advanced, composable visualizations that cater to the intricate relationships inherent in high-dimensional data. These capabilities are particularly important in the context of omics research, where the integration of diverse biological layers such as genomics, transcriptomics, epigenomics, and microbiome data presents significant visualization challenges.

A key strength of ggalign lies in its ability to offer flexible and dynamic layout options, including *StackLayout, QuadLayout*, and *CircleLayout* **(Figure 1 and Figure S1)**. These layouts enable researchers to select the most suitable format based on the nature of their data and the comparisons they wish to make. Importantly, ggalign eliminates the need for learning various syntaxes from different software or packages, providing an intuitive and cohesive interface for multi-plot data visualization, all within the familiar ggplot2 ecosystem.

Many existing solutions for composable visualization are restricted to one-to-one relationship, making them less adaptable for visualizing interconnected datasets **(Table 1)**. In contrast, ggalign offers a broader and more flexible approach, allowing users to visualize one-to-many, many-to-one, and many-to-many relationships, thus offering a more accurate reflection of the multi-dimensional nature of biological data. One of the distinguishing features of ggalign is its enhanced annotation capabilities, which facilitate clearer and more informative visualizations. By allowing the annotation of specific observations and datasets, ggalign significantly improves the interpretability of complex biological data. Our demonstrations using microbiome, transcriptomic, genomic, and single-cell gene expression data highlight how ggalign facilitates the integration of different biological layers, offering insightful visualizations that enable researchers to uncover deeper insights into complex biological phenomena **(Figure 2-3)**.

In conclusion, ggalign offers a powerful, flexible, and user-friendly solution for visualizing multi-dimensional biological data. Its ability to create aligned, composable visualizations that support both discrete and continuous variables makes it a valuable tool in the researcher’s toolkit, especially in the context of complex omics research. By facilitating clearer and more effective communication of complex data, ggalign significantly enhances the ability to interpret and extract meaningful insights from intricate biological datasets.

## Methods

### Dataset Collection

Pan-cancer data for the intratumor microbiome was obtained from the The Cancer Microbiome Atlas database (https://tcma.pratt.duke.edu/)^6^, while pan-cancer single-cell data was retrieved from Gene Expression Omnibus (https://www.ncbi.nlm.nih.gov/geo/)^7^ (accession: GSE210347^8^). Additionally, mutation data for PRAD was downloaded from The Cancer Genome Atlas (TCGA) (https://portal.gdc.cancer.gov/)^9,10^. Arm-level alterations of TCGA cancer data were obtained from the supplementary file of the study by Taylor et al.^11^. Data for Figure S2A was sourced from the supplementary file in the study by Wang X et al.^12^, and data for Figure S2B was obtained from https://github.com/YuLab-SMU/plotting-tree-with-data-using-ggtreeExtra^13^. Data for Figure 2A was downloaded from the supplementary file of the study by Thorsson et al.^5^, however, since the authors did not provide the clustering results, we chose to annotate only the representative immune signature, resulting in a slightly altered illustration from the original.

### Single-Cell Analysis

Gene expression matrices were normalized using the normalizeCounts function. To capture high cell-to-cell variation, 2,000 genes were selected based on their variance using the modelGeneVariances function. For Uniform Manifold Approximation and Projection (UMAP), we applied Mutual Nearest Neighbors integration^14^ to combine cells from different datasets, facilitating their joint analysis. The batch-corrected expression matrix of all integrated cells was then used to compute the UMAP. All analyses were conducted using the scrapper^15^ package.

### Statistics and reproducibility

Hierarchical clustering was performed using the *hclust()* function. The relative abundance of the intratumor microbiome was compared using the wilcoxon signed-rank test, with p-values adjusted using the Benjamini-Hochberg (BH) method. All analyses were conducted using the R stats package^16^.

## Data availability

All code to reproduce the analyses conducted in this study is publicly available on https://github.com/Yunuuuu/ggalign-research. For details regarding the data, please refer to the Methods section.

## Code availability

ggalign is open source and freely available on CRAN (https://CRAN.R-project.org/package=ggalign) and GitHub (https://github.com/Yunuuuu/ggalign/) under the MIT license. The official documentation website can be accessed at https://yunuuuu.github.io/ggalign-book/, and the full reference for the ggalign package is available at https://yunuuuu.github.io/ggalign/.

## Acknowledgements

We are grateful for resources from the Bioinformatics Platform, Furong Laboratory and Bioinformatics Center, Xiangya Hospital, Central South University. This work was funded by the National Natural Science Foundation of China (Grant Nos. 82303953, xxx), Central South University Startup Funding.

